# Microglia-dependent presynaptic disruption in an organotypic hippocampal slice culture model of neuroinflammation

**DOI:** 10.1101/502559

**Authors:** Oliva Sheppard, Michael P. Coleman, Claire S. Durrant

**Affiliations:** John van Geest Centre for Brain Repair, Department of Clinical Neurosciences, University of Cambridge, E.D Adrian Building, Forvie Site, Robinson Way Cambridge, CB2 OPY UK; Signalling Programme, Babraham Institute, Babraham Research Campus, Cambridge, CB22 3AT, UK

**Keywords:** Organotypic hippocampal slice culture, Lipopolysaccharide, Microglia, IL1β, Synapse, Synaptophysin, Presynaptic, Alzheimer’s disease

## Abstract

**Background:** Systemic inflammation, such as occurs during sepsis, bone fracture, infections or post-operative trauma, has been linked to synapse loss and cognitive decline in human patients and animal models. Organotypic hippocampal slice cultures (OHSCs) represent an underused tool in neuroinflammation; retaining much of the neuronal architecture, synaptic connections and diversity of cell types present in the hippocampus *in vivo* whilst providing convenient access to manipulate and sample the culture medium and observe cellular reactions as in other *in vitro* methods. Here we report the development of an OHSC model of synaptic disruption after aseptic inflammation and investigate the underlying mechanism.

**Methods:** OHSCs were generated from P6-P9 C57BL/6, the APP transgenic TgCRND8 model, or wild-type littermate mice according to the interface method. Aseptic inflammation was induced via addition of lipopolysaccharide (LPS) and cultures were analysed for changes in synaptic proteins via western blot. qPCR and ELISA analysis of the slice tissue and culture medium respectively determined changes in gene expression and protein secretion. Microglia were selectively depleted using the toxin clodronate and the effect of IL1β was assessed using a specific neutralising monoclonal antibody.

**Results:** Addition of LPS caused a loss of the presynaptic protein synaptophysin via a mechanism dependent on microglia and involving IL1β. Washout of LPS via medium exchange allows for partial recovery of synaptic protein levels after 2 weeks. TgCRND8 OHSCs do not show alterations in IL1β expression at a timepoint where they exhibit spontaneous synaptophysin loss, and LPS does not alter levels of APP or Aβ in wild-type OHSCs. This indicates that although synaptophysin loss is seen in both systems, there is likely to be distinct underlying pathogenic mechanisms between the neuroinflammatory and amyloid models.

**Conclusions:** We report the development of an OHSC model of LPS-induced synaptophysin loss and demonstrate a key role for microglia and involvement of IL1β. We propose that distinct molecular mechanisms lead to synaptophysin protein loss in LPS-exposed versus TgCRND8 OHSCs and provide a new experimental paradigm for assessing chronic changes in synaptic proteins, and synaptic plasticity, following acute inflammatory insults.

## Background

There is mounting evidence for a link between neuroinflammation, synapse loss and cognitive decline both in humans and in pre-clinical models. Systemic inflammatory events such as sepsis, periodontitis, infections, bone fracture and post-operative trauma can result in sustained high levels of circulating pro-inflammatory cytokines [1, 2] and have been linked with long-term cognitive impairment in patients [3–7]. In addition to seeding cognitive decline in previously healthy patients, systemic inflammation has also been shown to exacerbate neurodegenerative disease processes, being strongly associated with worse clinical outcome in multiple sclerosis [8, 9], accelerating cognitive decline in Alzheimer’s disease [2, 10–12] and inducing subacute motor deterioration associated with delirium in Parkinson’s disease [13].

In pre-clinical models, investigating the link between systemic inflammation and changes in the central nervous system (CNS) can be achieved by inducing aseptic inflammation using bacterial lipopolysaccharide (LPS), a potent endotoxin found on the cell walls of gram negative bacteria [14] which is seen to be elevated in plasma samples from septic patients [15, 16]. In animal models, LPS treatment has been shown to induce deficits in spatial learning assessed using the Morris water maze, as well a decline in the presynaptic protein synaptophysin in the hippocampus [17, 18]. In agreement with observations in human patients, it is often reported that inflammatory events worsen neurodegenerative disease processes in animal models. Notably, LPS injection has been shown to increase accumulation of intracellular APP and Aβ in APP_SWE_ mice [19], increase tau phosphorylation in rTg4510 mice [20] and accelerate motor and cognitive phenotypes in a mouse model of prion disease [21]. The ability of LPS to induce such changes in the CNS is believed to be due to its impact on the cytokine release profile of microglia. *In vitro* experiments have shown that murine neuronal cultures exposed to conditioned medium from LPS-treated microglia undergo a reduction in synaptic connections, likely due to an increase in IL1β production [22], whilst the ability of non-contact co-cultured microglia to induce synapse formation via IL-10 release is inhibited upon LPS application [23].

Whilst there has been much progress in understanding the links between synaptic changes and inflammatory stimuli, progress in this field would be facilitated by a model system that is as amenable to environmental/ pharmacological manipulation as *in vitro* models but also retains all the relevant cell population and synaptic architecture found *in vivo*. Organotypic hippocampal slice cultures, where thin sections of hippocampus are maintained in culture for several weeks [24–26], represent an excellent compromise between *in vivo* and *in vitro* models and have been surprisingly underused in the field of neuroinflammation. Previous studies examining the effect of LPS in OHSCs have focussed primarily on cell death and cytokine production, with LPS shown to prime cultures to subsequent insults such as AMPA-induced toxicity [27] or oxygen-glucose deprivation [28] as well as altering the production of pro-inflammatory cytokines [29] and growth factors [30]. Whilst synaptic disruption is commonly reported for both *in vitro* and *in vivo* LPS assays, there is currently only one publication examining these changes in OHSCs, which focussed on postsynaptic responses. It reported that application of high (1µg/ml) concentrations of LPS reduced the number of thin dendritic spines in CA1, resulting in a decreased frequency of excitatory post synaptic currents (EPSCs) [31]. As synaptic changes are likely to be clinically relevant to the cognitive decline seen in sepsis, or acceleration of neurodegenerative disease in human patients, there is great potential in expanding the OHSC model to further probe mechanisms of synaptic disruption, in particular presynaptic events, in response to inflammation.

In this study, we demonstrate that LPS treatment of OHSCs results in the depletion of the presynaptic protein synaptophysin in a manner dependent on microglia and involving IL1β. This differs from the molecular changes seen in an OHSC model of amyloid pathology using tissue from TgCRND8 mice, indicating that synaptophysin loss can be induced by distinct pathogenic mechanisms. We also assess the ability of OHSCs to recover after a transient inflammatory insult and explore whether the presynaptic disruption previously reported in TgCRND8 cultures [25] renders them more vulnerable to additional inflammatory insults.

## Methods

### Mice

Wild-type animals (C57BL/6Babr) were obtained from the breeding colony at the Babraham Institute. TgCRND8 mice [32] (which overexpress human APP695 with Swedish (K670N/M671L) and Indiana (V717F) mutations) were maintained as heterozygotes on a 62.5:37.5 SV129:C57BL/6 background, producing both transgenic and wild-type littermates. Animal work was approved by the Babraham Institute Animal Welfare, Experimentation and Ethics Committee and was performed in accordance with the Animals (Scientific Procedures) Act 1986 under Project License PPL 70/7620 and P98A03BF9. All animals were bred in a specific pathogen free animal facility with strict temperature and humidity control. Both genders were used in experiments.

### Organotypic Hippocampal Slice Cultures

OHSCs were cultured according to the interface method as described previously [24, 25]. Briefly, P6-P9 mouse pups were humanely sacrificed by cervical dislocation and their brains rapidly transferred to ice-cold dissection medium (EBSS + 25mM HEPES + 1x Penicillin/Streptomycin). Brains were transected at the midline, and glued (Loctite) to a vibratome stage. 350µm sagittal slices were cut using a Leica VT1000S vibratome and the hippocampus and associated entorhinal cortex dissected out. Slices were plated on 0.4μm pore membranes (Millipore: PICM0RG50) sitting on top of 1ml of maintenance medium (50 % MEM with Glutamax-1 (Life Tech:42360–024), 25 % Heat-inactivated horse serum (Life Tech: 26050–070), 23 % EBSS (Life Tech: 24010–043), 0.65 % D-Glucose (Sigma:G8270), 2 % Penicillin-Streptomycin (Life Tech: 15140–122) and 6 units/ml Nystatin (Sigma: N1638). 2–4 culture dishes per pup were made, depending on experimental protocol, with 2–3 slices plated per dish. 1 and 4 days after plating, cultures underwent a 100% medium exchange, before moving to a 50% weekly exchange thereafter. Cultures were maintained in an incubator under high humidity at 37°C and 5% CO_2_ for up to 5 weeks.

### Treatments

At 2 weeks *in vitro*, OHSCs were treated with 200ng/ml lipopolysaccharide from *E.Coli* O55:B5 (LPS) (Sigma L5418) or 20ng/ml murine Interleukin-1β (IL-1β) (Sigma I9401) for an additional 7 days. For microglial-depletion experiments, OHSCs were pre-treated with 100µg/ml clodronate (VWR: 233183) for 24 hours prior, and throughout, LPS treatment. For IL1β neutralising experiments, OHSCs were pre-treated with either a murine-IL1β neutralising mouse monoclonal antibody (Invivogen: mabg-mil1b) or a mouse IgG isotype control antibody specific to *E.Coli* β-Galactosidase (Invivogen: mabg1-ctrlm) for 24 hours prior, and throughout LPS treatment.

### Western Blotting

OHSCs were scraped off the membrane into ice-cold RIPA buffer (50mM Tris-HCl, 500mM NaCl, 1% Triton-X, 10nM EDTA, pH 8.0) with protease and phosphatase inhibitors (Thermofisher Scientific: 78442). Slices underwent probe sonication for 2 × 5 seconds to completely homogenise the tissue. Equal amounts of protein were denatured in Laemelli buffer (with 2-Mercaptoethanol) and loaded into 4–20% Tris-glycine gels for separation by SDS-PAGE. Proteins were transferred onto PDVF-FL prior to blocking in Odyssey blocking buffer for 1 hour at room temperature. Primary antibodies were diluted in 5% BSA in PBS-T with 0.05% sodium azide and membranes were incubated overnight at 4°C on the shaker. After 3 PBS-T washes, membranes were incubated in 1:10,000 secondary IRDye anti-mouse and anti-rabbit antibodies (Li-Cor) for 2 hours (protected from light), washed with PBS-T then imaged using a Li-Cor Odyssey CLX system. Band intensities were normalised to beta iii tubulin (Tuj1) to control for differences in neuron number. Primary antibodies were used as follows: 1:1,000 mouse synaptophysin (Abcam: ab8049), 1:500 rabbit PSD95 (Abcam: ab18258), 1:2,500 rabbit Tuj1 (Sigma: T2200).

### Immunohistochemistry

Slices remained adhered to membranes while fixed for 20 minutes in 4% PFA and then washed 3 times in PBS. The membranes were then cut and slices were placed in a 24-well plate and blocked for 1 hour in blocking solution (PBS + 0.5 % Triton X-100 + 3 % Goat Serum). Slices were incubated with primary antibody (1:500 Iba-1 (Alpha Laboratories: 019–19741) diluted in blocking solution overnight with shaking at 4°C. After 3 PBS washes, OHSCs were incubated (2 hours, room temperature, protected from light) with Alexa488 or 568 conjugated secondary antibodies (Life Technologies) diluted 1:250 in blocking solution. Slices were counterstained with Hoechst (1:5000 in PBS), washed in PBS then mounted on slides to be imaged via confocal microscopy.

### Quantitative PCR

RNA was extracted from OHSCs using the RNEasy Extraction Kit (Qiagen: 74104). From this, cDNA was synthesized using a Reverse Transcriptase Kit (Quantitect: 205310). Quantitative PCR was carried out using Brilliant III Ultra-Fast SYBR Green QPCR Master Mix (Agilent Technologies: 600882). The following PCR program was used on a BIO-RAD CFX96 Real-Time PCR Detection System with c1000 Touch Thermal cycler: 3 minutes at 95°C, 40 cycles of 5 seconds at 95°C, and 5 seconds minute at 60°C. Primers for each gene are listed in the table below.

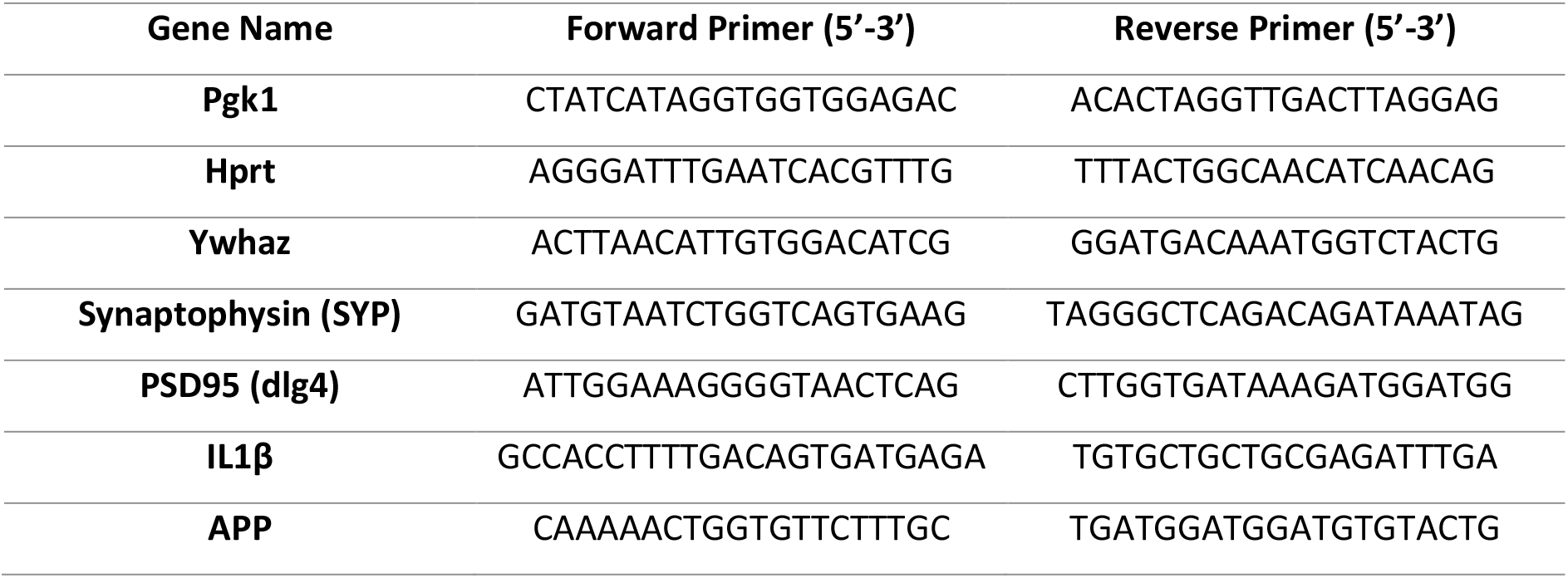

### ELISA

To determine the level of murine Aβ_1–42_ or IL1β in the culture medium, ELISAs were carried out using commercially available kits (Invitrogen: KMB3441 or R&D Systems: MLB00C). Medium was collected from slice cultures at various timepoints throughout LPS or IL-1β treatment. ELISA was carried out as per manufacturer’s instructions, with absorbance read using a PheraStar FS plate reader.

### Statistical Analysis

Data was analysed using GraphPad Prism Software. Statistical tests were chosen to match the data set type, including paired and un-paired T-tests and two way ANOVA. Significance values are reported as follows: p<0.05= *, p<0.01**, p<0.001***, p<0.0001***.

## Results

### LPS treatment induces the loss of the presynaptic protein synaptophysin

OHSCs were created from P6-P9 wild-type mice such that two separate culture dishes (each with 3 hippocampal slices) were generated per animal. Cultures were aged for 14 days *in vitro* before treatment with 200ng/ml LPS. Slices were collected for western blot or qPCR analysis after 7 days of treatment. For all analysis, LPS treated cultures were directly compared to the untreated control from the same animal. **Fig 1a** shows a representative western blot where lysates were probed for the presynaptic protein synaptophysin (SYP), postsynaptic protein PSD95 and neuronal microtubule protein beta-iii tubulin (Tuj1). Synaptic protein levels were normalised to Tuj1, to control for any loss of neurons that may confound any specific vulnerability of the synapses. LPS treatment resulted in a significant loss of synaptophysin (p=0.01) **(Fig 1b)** but did not alter the levels of PSD95 (p=0.67) **(Fig1c)**. qPCR analysis revealed that both synaptophysin **(Fig 1d)** and PSD95 (p=0.037) **(Fig 1e)** mRNA transcripts were reduced, although not significantly (p=0.058) for synaptophysin, in LPS treated cultures when compared to 3 housekeeping genes (Pgk1, Ywhaz and Hprt).

**Figure 1).**
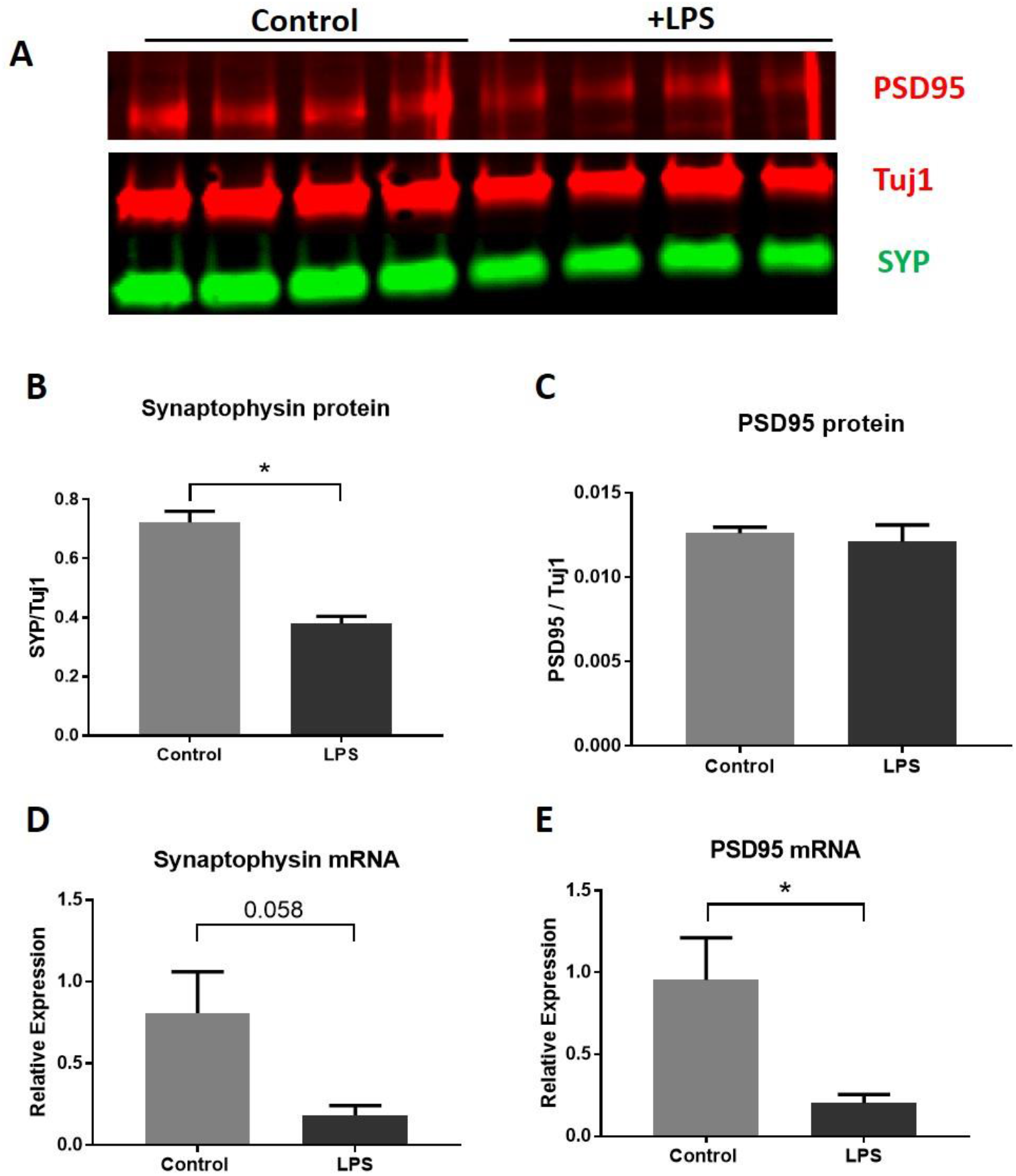
LPS addition causes reduction of synaptophsyin protein with no change in PSD95. 14 *days in vitro* organotypic hippocampal slice cultures were challenged with 200ng/ml LPS for a further 7 days. Slice cultures were harvested and synaptic proteins examined by western blot **(a)**. 7 days of LPS treatment result in loss of the presynaptic protein synaptophysin (p=0.0104*) **(b)** but with no change in the postsynaptic protein PSD95 (p=0.67) **(c)**. There is a strong trend for a reduction in synaptophysin mRNA (p=0.058) **(d)** and a significant decrease in PSD95 mRNA (p= 0.037*) **(e)**. All statistics were conducted using a paired t-test to account for matched control and treated OHSCs from the same animal.

### LPS-induced synaptophysin loss is microglial dependent

To test whether the loss of synaptophysin protein in LPS-treated OHSCs was due to alterations in microglia phenotype, microglia were depleted from slice cultures using the specific toxin clodronate [33–35] at 13 days *in vitro*. 24 hours later, half of the cultures received a further treatment of 200ng/ml LPS resulting in four treatment groups: control (no treatments), LPS only, clodronate only, and clodronate + LPS. Cultures were harvested at 21 days *in vitro* (7 days after LPS treatment). Immunofluorescence staining for the microglial marker Iba1 revealed a change in morphology and increase in number of microglia after LPS addition **(Fig 2a,b)**. Whilst the microglia detected in untreated (control) OHSCs were in a ramified, branched state **(Fig 2a)**, LPS treatment (in the absence of clodronate) resulted in an increase in total area of Iba1 signal as well as a shift to an amoeboid morphology **(Fig 2b)**. Pre-treatment with the microglial toxin clodronate resulted in almost complete depletion of microglia, even in the presence of LPS **(Fig 2c,d)**. The effect of microglial depletion on LPS-induced synaptophysin loss was assessed by western blot **(Fig 2e)** with the results showing that pre-treatment with clodronate before LPS addition could block the loss of synaptophysin protein **(Fig 2f)**. Whilst OHSCs treated with LPS in the absence of clodronate showed a significant reduction in synaptophysin (p=0.03), there was no difference between clodronate treated and clodronate + LPS treated synaptophysin levels (p=0.47). There was a significant rescue of synaptophysin levels when comparing LPS treated with clodronate + LPS treated cultures (p<0.001) This rescue indicates a role of OHSC microglia in the effect of LPS on presynaptic proteins. It is interesting to note that as well as preventing the LPS-induced synaptophysin loss, the addition of clodronate regardless of LPS treatment increase the levels of synaptophysin protein (two way ANOVA effect of clodronate P<0.0001) potentially indicating a role of microglia for reducing basal levels of synaptic protein in our OHSC model.

**Figure 2).**
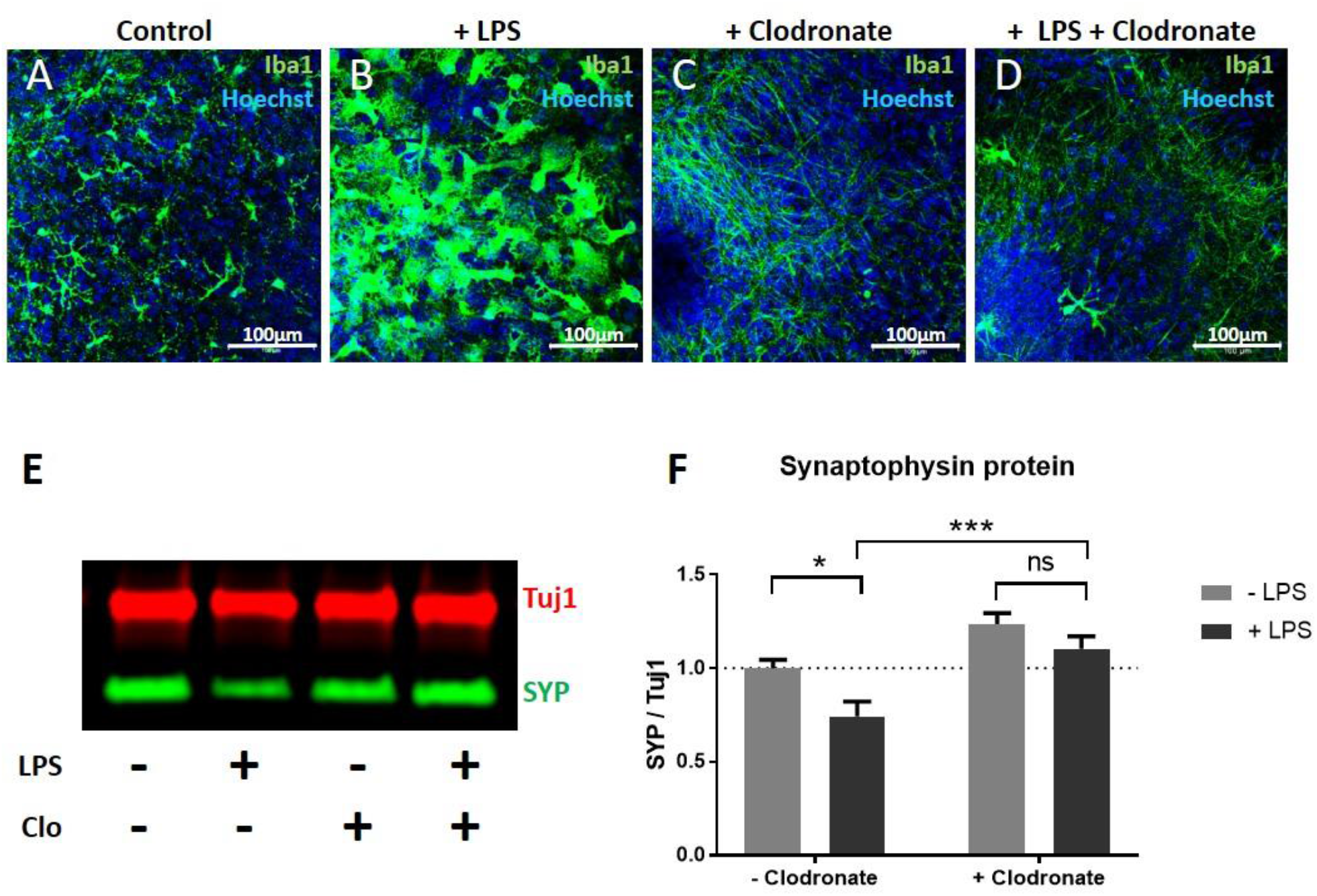
Pre-treatment with clodronate selectively kills microglia and prevents LPS-induced synaptophsyin loss. **(a-d)** Immunostaining of LPS and clodronate treated OHSCs (Green= Iba1, Blue= Hoechst). Whilst microglia in control conditions appear ramified **(a)** addition of LPS results in a striking alteration of microglial phenotype to an amoeboid morphology **(b)**. Pre-treatment of OHSCs with clodronate significantly reduces the number of microglia in LPS-naïve cultures **(c)** and LPS-exposed cultures **(d)**. Western blot of LPS and clodronate treated cultures **(e,f)** shows that whilst clodronate-naïve cultures show a reduction in synaptophysin when treated with LPS (p=0.03*) there is no difference between clodronate pre-treated cultures upon additional LPS application (p=0.47). There is a significant rescue seen when comparing the effect of clodronate pre-treatment on cultures treated with LPS (p=0.0009***) There is a significant overall effect of clodronate treatment regardless of LPS addition (p=<0.0001****). All statistics were conducted using a two-way ANOVA.

### IL1β is increased by LPS and is sufficient to induce synaptophysin depletion in OHSCs

To determine whether the LPS-induced loss of synaptophysin protein could be due to the release of inflammatory cytokines, the concentration of IL1β in the OHSC medium was determined by ELISA. Whilst in untreated culture medium, the levels of IL1β were undetectable, in LPS treated cultures there was an average of 6pg/ml IL1β detected, representing a significant increase (p=0.0008) **(Fig 3a)**. This observation occurs alongside a significant increase in IL1β mRNA in the slice tissue (p=0.016) **(Fig 3b)** indicating increased transcription of this inflammatory cytokine. To test whether the loss of synaptophysin protein seen after LPS addition could be due to the downstream production of IL1β, murine IL1β protein was applied directly to OHSCs at 14 days *in vitro*. After 7 days of 20ng/ml IL1β, OHSCs were harvested for western blot analysis **(Fig 3c)** As with LPS treatment (see **Fig 1** and **Fig 2**), there is a significant decrease in synaptophysin relative to Tuj1 (p=0.0029) **(Fig 3d)** and no significant change in PSD95 (p=0.16) **(Fig 3e)**. This demonstrates that IL1β application is *sufficient* to induce synaptophysin loss in OHSCs.

**Figure 3).**
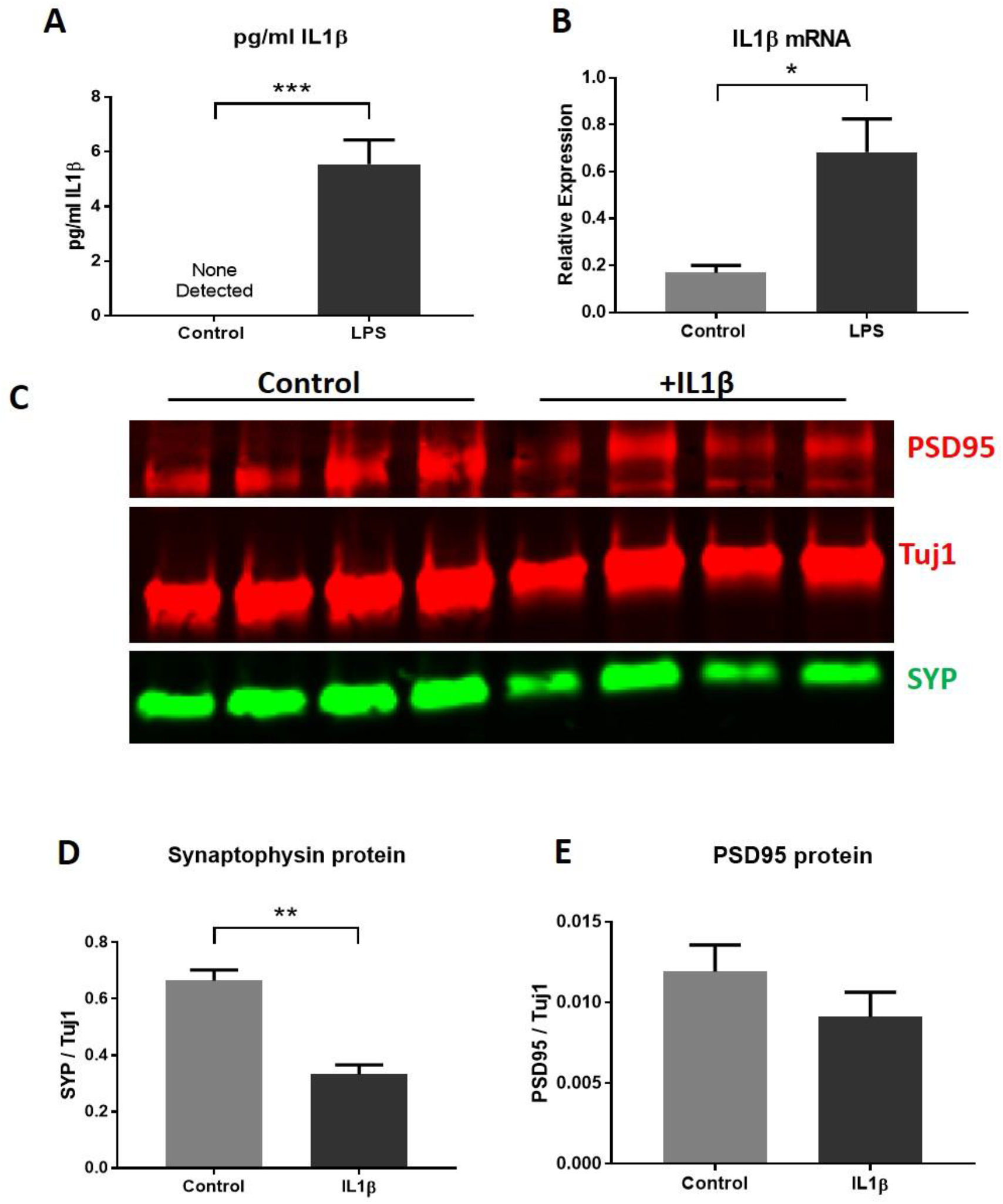
IL1β production is induced by LPS and addition of IL1β results in synaptophysin loss. LPS treated OHSCs show increased IL1β levels in the culture medium after 24 hours (p=0.0008) **(a)**. IL1β mRNA is also elevated in 3 weeks *in vitro* wild-type cultures treated with LPS for the last week *in vitro* (p=0.016) **(b)**. Treatment with 20ng/ml IL1β **(c)** results in reduced synaptophysin protein (p=0.0029) **(d)** with no significant change in PSD95 (p=0.16) **(e)**. All statistics were conducted using a paired t-test to account for matched control and treated OHSCs from the same anima

To determine whether IL1β is *necessary* for LPS to induce synaptophysin loss, 13 days *in vitro* OHSCs were treated with either a murine-IL1β neutralising mouse monoclonal antibody (α-IL1β) or a mouse IgG isotype control antibody specific to *E.Coli* β-Galactosidase (α-βGAL). 24 hours later, cultures were treated with 200ng/ml LPS. OHSCs were prepared such that the 4 different treatment conditions could be compared in tissue from the same animal, with synaptic protein levels in treated conditions compared relative to their tissue-matched LPS naïve α-βGAL control. Slices were harvested at 21 days *in vitro* and analysed by western blot **(Fig 4a)**. Whilst OHSCs treated with the isotype control antibody showed the expected loss of synaptophysin protein in response to LPS treatment (p=0.047), cultures treated with IL1β-neutralising antibody were not sensitive to the addition of LPS (p=0.52) **(Fig 4b)**. There was a non-significant, but trending, improvement in synaptophysin levels in IL1β-neutralising antibody +LPS OHSCs when compared to control antibody +LPS treated OHSCs (p=0.14). Therefore, while IL1β inhibition appears to partially rescue the LPS induced deficit in synaptophysin, a combination of factors, such as alternative inflammatory cytokines, are likely to be involved and may require further investigation.

**Figure 4).**
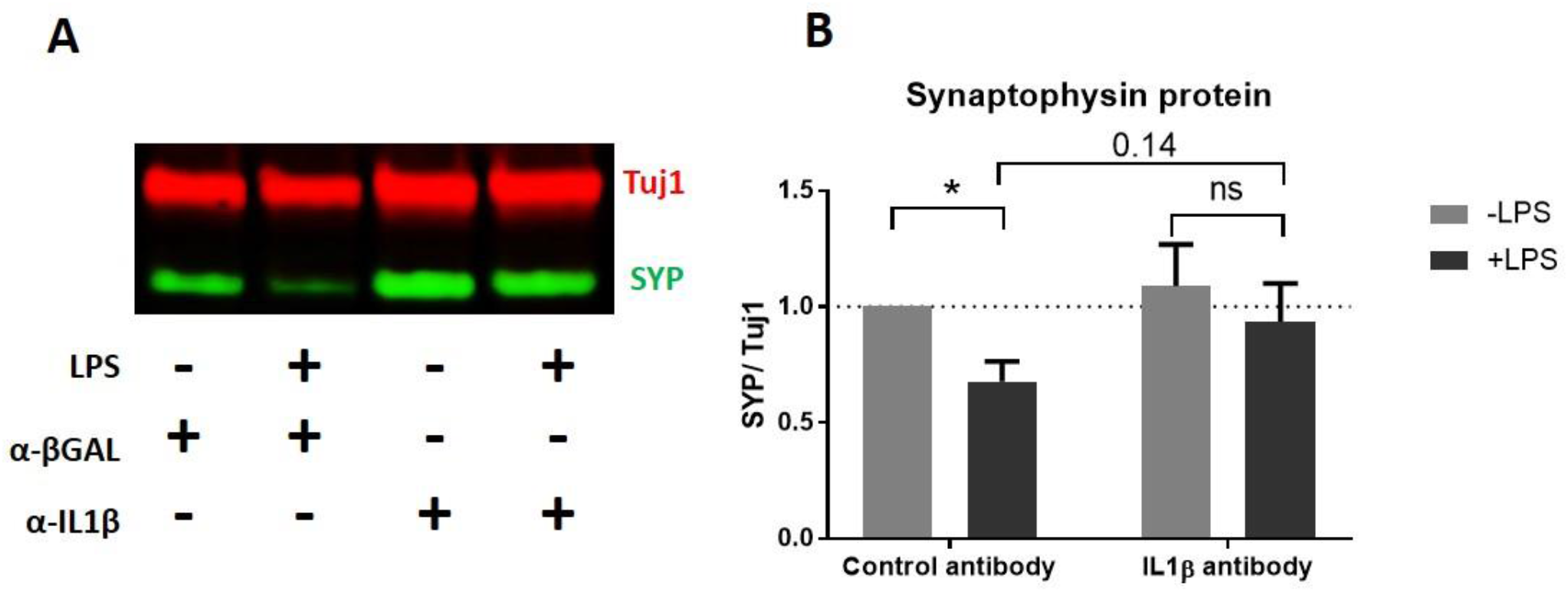
Application of anti-IL1β neutralising antibody alters the OHSC response to LPS. Western blot of antibody and LPS treated cultures **(a)** shows that whilst OHSCs pre-treated with anti-βGAL (control) antibody show a reduction in synaptophysin when treated with LPS (p=0.047*) cultures pre-treated with anti-IL1β neutralising antibody are resistant to LPS addition (p=0.52) **(b)**. There is a non significant trend for improvement of synaptophysin levels when comparing control antibody +LPS and anti-IL1β antibody +LPS (p=0.135). All analysis was conducted using a two way ANOVA with samples normalised to their tissue-matched Control antibody -LPS control.

### Assessing synaptic protein recovery after LPS removal

A key advantage of the slice culture system is the ability to rapidly and accurately manipulate the extracellular environment in a way that is exceedingly difficult to achieve *in vivo*. We sought to determine whether complete removal of the inflammatory stimulus (LPS) after the depletion of synaptophysin has already taken place permits for the recovery of synaptic protein to un-treated levels. OHSCs were prepared such that 2 culture dishes were created per mouse (each with 3 slices). After 2 weeks *in vitro* all of the cultures underwent a 100% medium exchange, with 1 of the 2 dishes receiving 200ng/ml LPS. After a further week of treatment, one group of slices (those representing a 0 week post-treatment timepoint) were harvested for western blot whilst all other cultures underwent a further 100% medium exchange, receiving untreated medium. Slices were then left to “recover” for a further 1 or 2 weeks *in vitro* **(Fig 5a)**. Synaptophysin protein levels in OHSC lysates were analysed by western blot **(Fig 5b)**. As expected, OHSCs harvested immediately after LPS treatment showed a reduction in synaptophysin protein when compared to untreated slices from the same animal (p=0.014) **(Fig 5c)**. At 1 week after LPS removal, synaptophysin levels were not significantly different in treated vs untreated controls (p=0.30) and, whilst there is a trend for a reduction at 2 weeks, this is not significant (p=0.13). This indicates that loss of synaptophysin in response to LPS is substantially reversible after the inflammatory insult is removed, although we cannot exclude the possibility of some limited longer-term effect at present.

**Figure 5).**
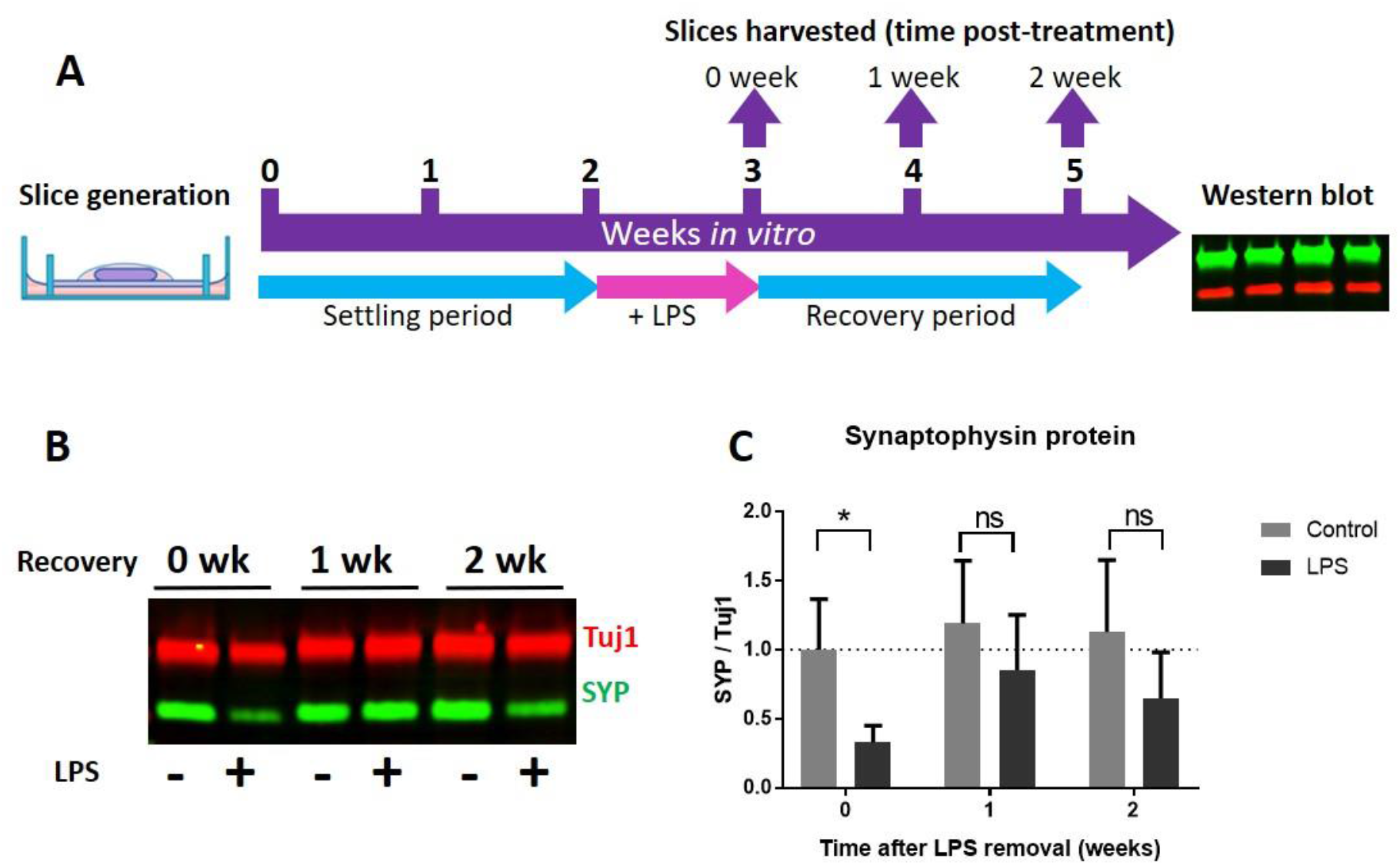
Synaptophysin protein levels undergo recovery after LPS removal. A diagrammatic representation of the LPS recovery experiment **(a)**. OHSCs are aged for 2 weeks *in vitro* before undergoing 1 week of 200ng/ml LPS. At 3 weeks *in vitro* some slices are harvested to represent a 0-weeks after LPS removal timepoint. All other cultures undergo a 100% medium exchange to LPS-free medium. Slices are then harvested at either 1 week or 2 weeks after LPS removal and synaptophysin protein levels assessed by western blot **(b)**. Slices harvested with no recovery after LPS removal showed a reduction in synaptophysin levels when compared to untreated samples (p=0.014). At 1 week (p=0.30) or 2 weeks (p=0.13) after LPS removal, there is no significance between LPS-exposed and untreated OHSCs. **(c)**. Analysis was conducted using a two way ANOVA.

### LPS induced synaptophysin loss does not work through an altered amyloid pathway in OHSCs

In our previous work, we reported a progressive loss of synaptophysin protein in OHSCs from the Alzheimer’s disease mouse model TgCRND8 [25]. As inflammation is often linked to increased cognitive decline in people living with dementia [10], we tested the hypothesis that the effects of LPS on wild-type OHSCs may act through mechanisms related to those in Alzheimer’s disease. The main features of the TgCRND8 model are overexpression of amyloid precursor protein (APP) and subsequent overproduction of Aβ. We saw no change in APP mRNA in wild-type OHSCs treated with LPS (p=0.20) **(Fig 6a)** and treatment with LPS (p=0.053) **(Fig 6b)** or IL1β (p=0.56) **(Fig 6c)** did not increase the production of murine Aβ_1–42_ (as measured by protein concentration in the culture medium). Indeed LPS treatment significantly *reduced* the detectable level of Aβ in the culture medium over time. We next sought to determine whether the previously reported synaptophysin loss seen in TgCRND8 OHSCs could coincide with an increased production of IL1β, as reported here for our LPS treated OHSCs **(Fig 3,4)**. There was no significant difference in IL1β mRNA levels in 5 weeks *in vitro* TgCRND8 OHSCs compared to wild-type littermate controls (p=0.37) **(Fig 6d)**, despite this being a timepoint where there is known to be synaptophysin depletion [25].

**Figure 6).**
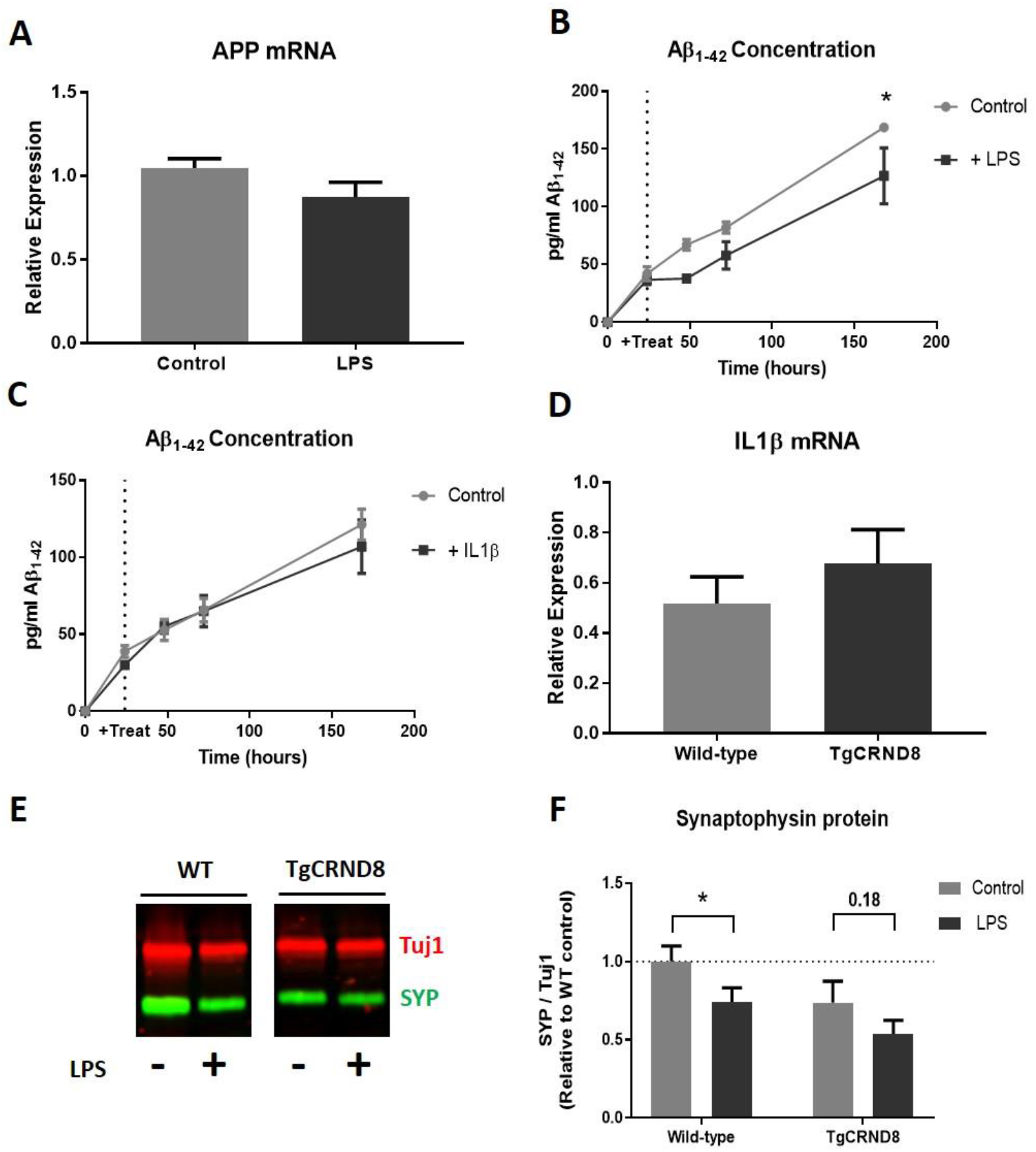
LPS does not interact with the amyloid pathway in OHSCs. LPS treatment does not alter APP mRNA expression levels (p=0.20)**(a)**. ELISA analysis of OHSC medium reveals that LPS treatment reduces the production of Aβ_1–42_ (Two way ANOVA: effect of treatment p=0.053) **(b)** whilst IL1β application does not influence Aβ_1–42_ accumulation (Two way ANOVA effect of treatment p=0.56) **(c)**. 5 week i*n vitro* TgCRND8 cultures do not show increased IL1β mRNA levels when compared to wild-type littermate controls (p=0.37) **(d)**. Representative western blot of 3 week *in vitro* wild-type and TgCRND8 OHSCs after 1 week of LPS **(e)**. There is a strong trend for TgCRND8 cultures to have lower synaptophysin protein levels regardless of LPS addition (two way ANOVA effect of genotype p=0.076). Whilst LPS addition to wild-type OHSCs reduces synaptophysin levels (p=0.027*), this was not significant in TgCRND8 cultures (p=0.18)

Whilst LPS treatment does not alter the amyloid pathway in wild-type OHSCs, it is important to test whether TgCRND8 OHSCs are more sensitive to LPS-induced synaptophysin loss. OHSCs were prepared from TgCRND8 and wild-type litter mates such that 2 cultures (with 3 slices per dish) were produced from each mouse. After 2 weeks *in vitro* one dish from each pair was treated with 200ng/ml LPS. At 3 weeks *in vitro* OHSCs were harvested and synaptophysin protein levels analysed by western blot **(Fig 6e)**. TgCRND8 cultures showed a strong trend for reduced synaptophysin levels when compared to wild-type OHSCs, regardless of LPS treatment (two way ANOVA p=0.076) **(Fig 6f)**. Interestingly, whilst wild-type OHSCs showed a reduction in synaptophysin in response to LPS (p=0.027), this was not significant in TgCRND8 cultures (p=0.18). This indicates that, under this experimental paradigm, TgCRND8 cultures do not show enhanced sensitivity to LPS, although the possibility that the effects can be additive remains to be explored.

## Discussion

In this work, we have shown that LPS addition to organotypic hippocampal slice cultures results in the depletion of the presynaptic protein synaptophysin in a mechanism dependent on microglia and involving IL1β activity. We have also provided a new model for assessing synaptic recovery after an inflammatory insult, which we envisage will be of great utility as a tool to probe key questions relating to the impact of acute inflammatory events on long term synaptic health and plasticity. By comparing the effect of LPS on wild-type and TgCRND8 cultures, we explored whether tissue that is already undergoing synaptic disruption is rendered more vulnerable to inflammatory insults and whether there is a common mechanism of action. In this experimental paradigm, we found that TgCRND8 and LPS-treated cultures do not show similarities in their molecular changes, namely APP expression, Aβ production and upregulation of IL1β. Whilst, perhaps surprisingly, TgCRND8 cultures did not show a greater loss of synaptic protein in response to LPS, our experiments raise interesting questions about the mechanisms of synaptophysin loss in Alzheimer’s-like and inflammatory conditions and provide an excellent platform to develop future studies of both mechanism and therapeutic strategies.

We have shown, for the first time in an OHSC model, that LPS treatment induces the loss of the presynaptic protein synaptophysin. This is in agreement with reports from *in vivo* and *in vitro* studies linking LPS administration with presynaptic protein depletion [17, 22] but, as demonstrated here, greatly facilitates mechanistic studies in the context of normal hippocampal circuits. Interestingly, a number of studies have shown strong correlations between the levels of synaptophysin and cognitive impairment after LPS exposure [17, 36, 37], identifying this as a clinically relevant pathogenic change. Whilst at the time of writing there are no studies examining the levels of synaptic protein in human brain after acute inflammatory insults, studies into Alzheimer’s disease [38–40], amyotrophic lateral sclerosis [41] and frontotemporal dementia [42] have all shown that the loss of synapses and synaptic protein is a strong correlate of clinical outcome. Understanding the mechanisms leading to synaptophysin depletion is therefore likely to be of great utility in identifying methods for treatment of patients at risk of cognitive decline after systemic inflammation.

Previous work looking at synaptic alterations in OHSCs after LPS application reported a loss of dendritic spines and associated reduction in EPSPs [31]. This post-synaptic deficit is in contrast to our data showing that there is no alteration in the level of PSD95 protein in our OHSC model. These results, however, are not conflicting, and could raise interesting questions about the nature of pre- and post-synaptic vulnerability. Of note, is their finding that there was a dose-dependent effect of LPS on dendritic spines, with the greatest loss seen using a dose 5 x higher than used in this study. The authors also do not report having examined the presynaptic compartment in their work, so it is feasible that both a pre- and post-synaptic deficit is present in their model. Together with our finding that despite differences in the extent of protein depletion, both synaptophysin and PSD95 show reductions in mRNA, this could imply that whilst both compartments are sensitive to disruption by LPS, presynaptic disruption, potentially partially at the level of transcription, occurs earlier and under lower levels of inflammatory insult. This would align with previous reports that changes in presynaptic proteins occur prior to changes to post-synaptic proteins [43–45]. The reduction in PSD95 transcript without an associated change in protein could be due to PSD95 having a lower turnover in OHSCs, rendering it more resistant to alterations. Likewise, the presence of PSD95 in a western blot does not prove that this protein exists in structurally normal spines so we cannot rule out changes to the organisation of the post-synaptic compartment. Future work could examine if PSD95 concentration begins to fall after a longer exposure time, or higher dose, of LPS and test whether both compartments are affected directly or whether the effect on one is secondary to the other.

Our finding that the LPS-dependent loss of synaptophysin in OHSCs is reliant on the presence of microglia agrees with a number of *in vitro* studies demonstrating that LPS-induced changes in microglial cytokine profile result in loss of synaptic proteins [22, 23] and that other supporting cells, such as astrocytes, do not respond to LPS as they lack the TLR4 receptor [46]. Most of these studies involve transfer of conditioned medium from LPS-treated microglia onto primary neuronal cultures which, although able to demonstrate that components released into the medium from microglia can be synaptotoxic, could be an over simplistic representation of the mechanisms at play in the CNS. The reaction of a primary cultured microglia *in vitro* may be different to microglia that are in an environment containing all the relevant CNS cell types in their normal cytoarchitecture. Cross talk between microglia, neurons and other supporting cells, including highly localised effects, is likely to be of key importance when examining how inflammatory insults can result in clinically relevant synaptic alterations [47]. As well as retaining the different cell types in a system that is more representative of the *in vivo* architecture, the amenability of OHSCs to pharmacological intervention allowed us to effectively and specifically deplete microglia using clodronate, something that would be difficult to achieve *in vivo*. This demonstrated that microglia are a vital component of LPS-induced synaptophysin loss in the OHSCs.

Our data demonstrating that LPS-induced IL1β production likely contributes to the loss of synaptophysin in OHSCs also agrees with previous reports *in vitro*, where primary neuronal cultures treated with conditioned medium from LPS-exposed microglia were protected from synapse loss via inhibition of IL1 receptors [22]. Here, we pre-treated OHSCs with a neutralising antibody against IL1β prior to LPS application and found that, whilst this prevented alterations to synaptophysin when compared to LPS naïve anti-IL1β antibody OHSCs, it only afforded partial rescue of the phenotype when compared to cultures exposed to a control antibody and LPS. This indicates that, whilst IL1β is sufficient to cause loss of synaptophysin and its inhibition may be therapeutically beneficial, future work will be required to determine if other LPS-induced factors (including a broader range of inflammatory cytokines) may need to be simultaneously targeted to completely prevent the loss of synaptophysin. In comparison to microglial depletion, the neutralisation of IL1β (or other cytokines) could be a more targeted method to protect synapses after an inflammatory insult in a clinical setting. As the production of IL1β increases after LPS addition, and can readily be detected by ELISA, it may be possible to test patient blood or CSF samples after a potential inflammatory insult to assess whether synaptic damage is likely to occur and treat accordingly. As our current work has only demonstrated efficacy of clodronate *prior* to LPS addition, further studies need to be carried out to determine at what point after an inflammatory insult inflammatory cytokines be neutralised to prevent loss of synaptic protein.

Our observation that complete removal of LPS after synaptic protein loss has taken place permits recovery approaching un-treated levels, is of great interest. Firstly, it demonstrates the utility of the slice culture system in probing mechanistic questions in this field, including mechanisms of synaptic plasticity. The ability to rapidly and accurately manipulate the extracellular environment by way of the culture medium provides an experimental advantage over animal models, whilst retaining some *in vivo*-like advantages such as a representative cell populations and neuronal architecture. Secondly, it provides an indication that at least partial recovery of function is possible if the inflammatory insult is removed. Whilst studies in human patients often report long term cognitive deficits or worsening of neurodegenerative disease processes after acute inflammatory insults [4], it is often seen that these patients retain high levels of circulating inflammatory cytokines [1]. Devising treatments that could “reset” the extracellular environment to a non-inflammatory state could be of benefit when seeking to prevent cognitive decline. Alternatively, it may be that the recovery of synaptic proteins can occur in some circumstances and fail in others. A study by Hao et al. found that treatment of pregnant rats with LPS resulted in their offspring showing significant neurodevelopmental brain damage, resulting in reduced synaptic protein and cognitive deficits that continue to worsen as the animal aged [48]. In this case, the retention of the synaptic deficit may be explained by the damage occurring during a critical time in development. Conversely, advanced age or already damaged synapses may be more vulnerable to lasting damage after an inflammatory insult. One limitation of OHSCs in this regard is the need to generate them from neonatal brains. However, we sought to examine part of this question by asking whether TgCRND8 cultures, which show progressive depletion of presynaptic proteins, would be more sensitive to LPS addition. Whilst we confirmed that the TgCRND8 OHSCs had a lower level of synaptophysin regardless of treatment, we did not see a significant decline in response to LPS. It could be that the synaptophysin loss was already at a maximal level potentially explaining the lack of significant additive effect. Future work could explore the difference between wild-type and TgCRND8 OHSCs in our recovery paradigm to answer the question of whether TgCRND8 slices (or transgenic OHSCs of interest) are less able to recover, and thus more susceptible to *chronic* or repeated damage as opposed to greater *acute* damage, after a single inflammatory insult.

Our finding that TgCRND8 slices do not overproduce IL1β at a time when they exhibit synaptophysin depletion, in addition to the absence of changes in the amyloid pathway upon inflammatory insults tested here, makes it likely that these different OHSC models represent different mechanisms of synaptic disruption. How systemic inflammation interacts with the amyloid pathway is a hotly debated topic, with studies demonstrating both *enhancement* [19, 49] and *reduction* [50, 51] of Aβ accumulation in response to LPS administration *in vivo*. Future work comparing how perturbations in the amyloid cascade or aseptic induction of neuroinflammation result in the loss of the same protein may help elucidate common or divergent pathways. Interestingly, virtually complete removal of microglia in an Alzheimer’s disease mouse model also did not alter the progression of cerebral amyloidosis, potentially providing further evidence for divergence in inflammatory and amyloidogenic pathogenesis that could be explored further in our slice culture system [52]. Targeting drug development to specific types of insult could help tailor treatments in patients, potentially affording greater efficacy with a reduction in unwanted side effects. The slice culture system represents an excellent tool to probe different mechanisms of presynaptic protein loss.

## Conclusions

In summary, we have shown that the addition of LPS to OHSCs results in loss of the presynaptic protein synaptophysin coinciding with increased expression of IL1β. The depletion of synaptic proteins can be prevented by pre-treatment with the microglia-specific toxin clodronate prior to LPS exposure. Application of an IL1β neutralising antibody prevents significant loss of synaptophsyin upon additional LPS application and shows a trend for rescue when compared to control slices exposed to LPS. By treating TgCRND8 and wild-type littermate OHSCs with LPS, we explored whether ongoing synaptic disruption in an OHSC model of amyloid pathology would increase vulnerability to inflammatory insults. Whilst, in this work, we did not find strong evidence for a significant additive effect of LPS when applied to TgCRND8 cultures, comparing LPS-exposed and amyloid transgenic OHSCs revealed differences in APP expression, Aβ generation and IL1β production that may indicate divergent pathogenic mechanisms leading to synaptophysin loss. We also report an experimental paradigm for assessing chronic synaptic changes after an acute inflammatory insult, demonstrating that LPS-induced loss of synaptophysin is a partially reversible change in our OHSC model.

**List of Abbreviations**
Aβ: Amyloid beta peptide
APP: Amyloid precursor protein
ELISA: Enzyme linked immunosorbent assay
IL1β: Interleukin 1β
LPS: Lipopolysaccharide
OHSC: Organotypic hippocampal slice culture
PBS: Phosphate buffered saline
PBS-T: Phosphate buffered saline with 0.1% Tween-20
PSD95: Post synaptic density 95
SYP: Synaptophysin

## Declarations

### Ethical approval and consent to participate

Animal work was approved by the Babraham Institute Animal Welfare, Experimentation and Ethics Committee and was performed in accordance with the Animals (Scientific Procedures) Act 1986 under Project License PPL 70/7620 and P98A03BF9.

### Consent for publication

Not applicable

### Availability of data and materials

The datasets used and/or analysed during the current study are available from the corresponding author on reasonable request.

## Author’s Contributions

Study concept and design: OS, MC and CD. Acquisition of data: OS and CD. Statistical Analysis: OS and CD. Analysis and interpretation of the data: OS, MC and CD. CD, OS and MC co-wrote the manuscript. All authors read and approved the final manuscript.

## Acknowledgements

We would like to thank the Babraham Biological Support Unit staff for their work involving the breeding and maintenance of mice used in this study.

## Author’s Information

Claire Durrant was previously known as Claire Harwell.

